# Immunometabolic cues recompose and reprogram the microenvironment around biomaterials

**DOI:** 10.1101/2023.07.30.551180

**Authors:** Chima V. Maduka, Axel D. Schmitter-Sánchez, Ashley V. Makela, Evran Ural, Katlin B. Stivers, Hunter Pope, Maxwell M. Kuhnert, Oluwatosin M. Habeeb, Anthony Tundo, Mohammed Alhaj, Artem Kiselev, Shoue Chen, Andrew J. Olive, Kurt D. Hankenson, Ramani Narayan, Sangbum Park, Jennifer H. Elisseeff, Christopher H. Contag

## Abstract

Circulating monocytes infiltrate and coordinate immune responses in various inflamed tissues, such as those surrounding implanted biomaterials, affecting therapeutic, diagnostic, tissue engineering and regenerative applications. Here, we show that immunometabolic cues in the biomaterial microenvironment govern CCR2- and CX3CR1-dependent trafficking of immune cells, including neutrophils and monocytes; ultimately, this affects the composition and activation states of macrophage and dendritic cell populations. Furthermore, immunometabolic cues around implants orchestrate the relative composition of proinflammatory, transitory and anti-inflammatory CCR2^+^, CX3CR1^+^ and CCR2^+^CX3CR1^+^ immune cell populations. Consequently, modifying immunometabolism by glycolytic inhibition drives a pro-regenerative microenvironment in part by myeloid cells around amorphous polylactide implants. In addition to, Arginase 1-expressing myeloid cells, T helper 2 cells and γδ^+^ T-cells producing IL-4 significantly contribute to shaping the metabolically reprogramed, pro-regenerative microenvironment around crystalline polylactide biomaterials. Taken together, we find that local metabolic states regulate inflammatory processes in the biomaterial microenvironment, with implications for translational medicine.

## Introduction

Control over the immune response to implanted biomaterials is required for medical devices to effectively function in therapeutic, diagnostic, tissue engineering and regenerative applications^1-4^. Trafficking of monocytes from the bloodstream into tissues is mediated by CCR2 signaling^5^, and is crucial to the foreign body response because extravasated monocytes develop into macrophages and dendritic cells, mediating the inflammatory response around implanted biomaterials. Recent advances in understanding metabolic adaptation in the biomaterial microenvironment reveal that changes in immune cell metabolism orchestrate immunological events in complex ways that could be leveraged to enhance regenerative medicine^6-13^. However, how local immunometabolic cues in the biomaterial microenvironment regulate the trafficking of immune cells and organize the relative composition of proinflammatory, transitory and anti-inflammatory or pro-regenerative phenotypes has not been elucidated. Despite a developing understanding of the changing phenotype(s) of recruited CCR2^+^ and CX3CR1^+^ cells in response to sterile injury and in several disease models^14-19^, the immunophenotypic composition of these cells in the biomaterial microenvironment has not been studied.

By fabricating amorphous polylactide (aPLA) implants, with and without embedded metabolic modulators, and locally implanting these biomaterial formulations in fluorescent reporter and wild-type mice, we demonstrate that the trafficking of immune cells to biomaterials is dependent on local metabolic cues. Additionally, prevailing immunometabolic cues in the biomaterial microenvironment regulate the relative composition of polarized CCR2^+^ and CX3CR1^+^ populations, with metabolic changes able to create a pro-regenerative phenotype with the help of myeloid cells in the aPLA biomaterial microenvironment. In addition to Arginase 1-expressing myeloid cells, T helper 2 cells and γδ^+^ T-cells producing IL-4 contribute to shaping the pro-regenerative microenvironment surrounding crystalline polylactide (cPLA).

### Rewiring metabolism in the amorphous polylactide biomaterial microenvironment controls CCR2- and CX3CR1-dependent trafficking

To understand the role of local metabolic cues in the trafficking of immune cells to sites of biomaterial implantation, we crossed *Ccr2*^*RFP/RFP*^*Cx3cr1*^*GFP/GFP*^ (CCR2- and CX3CR1-deficient) mice to B6 albino mice (Fig. 1a). The resulting F1 generation, *Ccr2*^*RFP/+*^*Cx3cr1*^*GFP/+*^ (CCR2- and CX3CR1-expressing) mice, were surgically incised (without biomaterial implantation) as sham controls; implanted with 7.5mm long reprocessed amorphous polylactide (hereafter, referred to as aPLA); or implanted with aPLA incorporating either 2-deoxyglucose (2DG) or aminooxyacetic acid (a.a.) at previously optimized concentrations^20^. In these dual reporter mice, RFP (CCR2) is expressed in ≈ 90% of Ly6C^+^ cells, allowing RFP to predominantly track classical monocytes, the analogue of CD14^+^ monocytes in humans^17,18^. GFP (CX3CR1) is mostly expressed by Ly6C^-^ alternative or resident monocytes, the analogue of CD16^+^ monocytes in humans^17^, as well as a subset of natural killer and dendritic cells^17,19^. Thus, intravital microscopic imaging of tissues adjacent to the implants allowed for visual monitoring of immune cells in the biomaterial microenvironment (Supplementary Fig. 1a). We used two inhibitors that act at different metabolic steps; 2DG inhibits hexokinase in the glycolytic pathway^21^, a.a. reduces both mitochondrial uptake of glycolytic substrates^22^ and glutamine metabolism^23^. Lastly, we included sham and aPLA groups among *Ccr2*^*RFP/RFP*^*Cx3cr1*^*GFP/GFP*^ mice (effectively knockout mice for these receptors; Supplementary Fig. 2a) to understand the contribution of CCR2- and CX3CR1-expression in the recruitment of immune cells to the biomaterial microenvironment.

**Figure 1.**
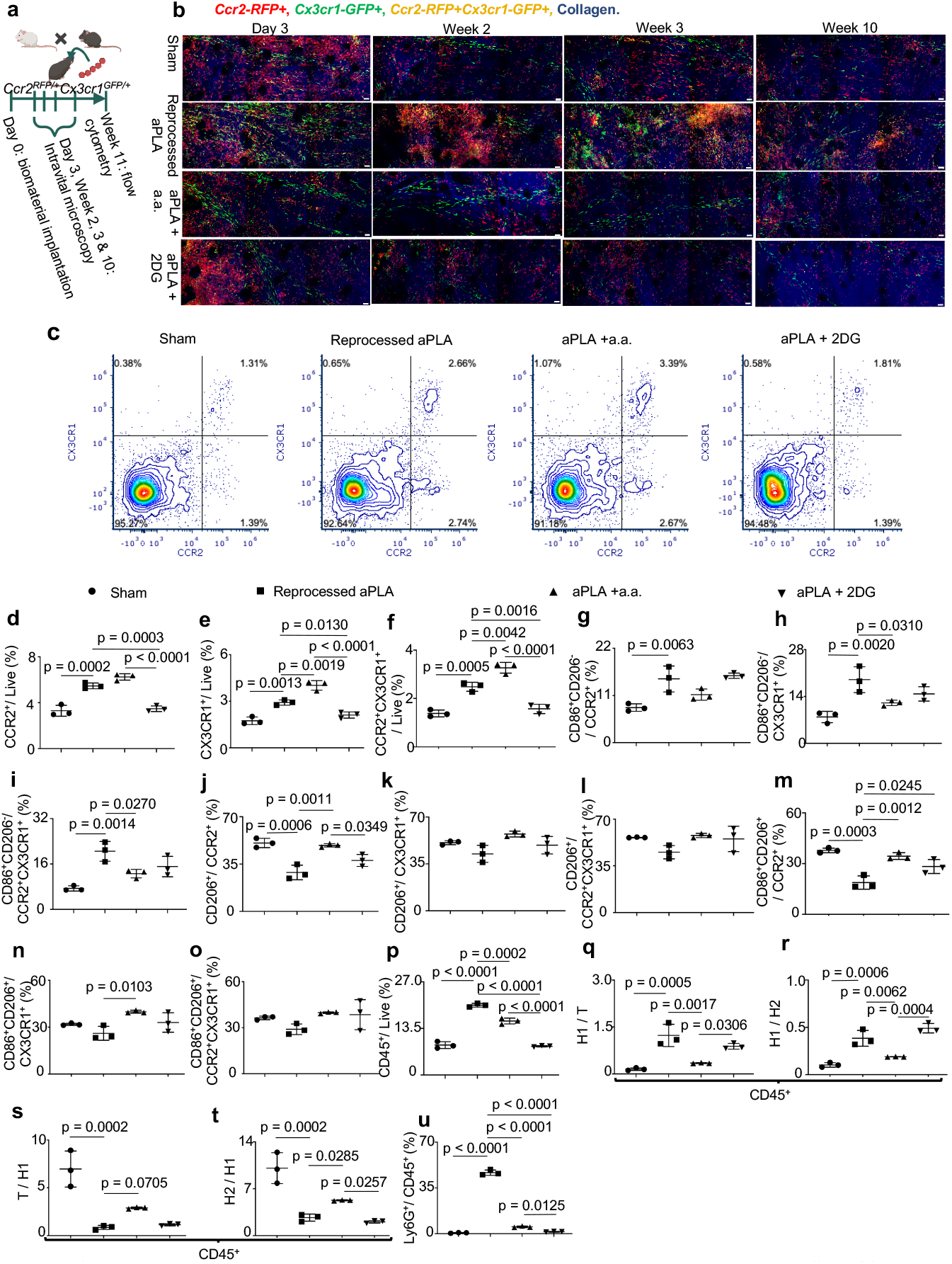
Locally rewiring immune cell metabolism in the amorphous polylactide biomaterial environment affects CCR2- and CX3CR1-dependent trafficking. **a**, B6 albino mice were crossed to *Ccr2*^*RFP/RFP*^*Cx3cr1*^*GFP/GFP*^ mice to generate *Ccr2*^*RFP/+*^*Cx3cr1*^*GFP/+*^ mice, which were surgically incised (sham group) or implanted with biomaterials. Intravital microscopy preceded flow cytometric analysis of tissues around incision sites (sham group) or biomaterials. **b**, Representative intravital microscopy images around incision sites (sham group), reprocessed amorphous polylactide (aPLA), aPLA incorporating aminooxyacetic acid (a.a.) or 2-deoxyglucose (2DG); scale bars are 50 μm. **c**, Representative CCR2 and CX3CR1 flow cytometry plots gated on live cells. **c-e**, Flow cytometry quantification of CCR2^+^(c), CX3CR1^+^(d) and CCR2^+^CX3CR1^+^(e) cells. **f-h**, Quantification of proinflammatory (CD86^+^CD206^-^) cells among CCR2^+^(f), CX3CR1^+^(g) and CCR2^+^CX3CR1^+^(h) populations. **i-k**, Quantification of anti-inflammatory (CD206^+^) cells among CCR2^+^(i), CX3CR1^+^(j) and CCR2^+^CX3CR1^+^(k) populations. **l-n**, Quantification of transition (CD86^+^CD206^+^) cells among CCR2^+^(l), CX3CR1^+^(m) and CCR2^+^CX3CR1^+^(n) populations. **o**, Nucleated hematopoietic (CD45^+^) cells. **p-q**, Fold change of proinflammatory (H1; CD86^+^CD206^-^) cells with respect to transition (T; CD86^+^CD206^+^) cells (p) or anti-inflammatory (H2; CD206^+^) cells (q). **r-s**, Fold change of T (r) or H2 (s) cells with respect to H1 cells. **t**, Neutrophils (CD45^+^Ly6G^+^ cells). One-way ANOVA followed by Tukey’s or Newman-Keul’s multiple comparison test, n = 3 mice per group.

We observed an initial increase in CCR2- and CX3CR1-expression that declined over time in all groups of *Ccr2*^*RFP/+*^*Cx3cr1*^*GFP/+*^ mice by intravital microscopy (Fig. 1b). CCR2^+^ and CX3CR1^+^ cells were observed in tissues of *Ccr2*^*RFP/+*^*Cx3cr1*^*GFP/+*^ mice, including sham (Supplementary Video 1), aPLA (Supplementary Video 2), aPLA + a.a. (Supplementary Video 3) and aPLA + 2DG (Supplementary Video 4) groups 10 weeks after surgeries. Notably, CCR2- and CX3CR1-expression appeared elevated in the aPLA group of *Ccr2*^*RFP/+*^*Cx3cr1*^*GFP/+*^ mice (Fig. 1b; Supplementary Video 2). Flow cytometric (quantitative) analyses of tissues around the implants corroborated increased CCR2^+^, CX3CR1^+^ and CCR2^+^CX3CR1^+^ cell populations in the aPLA group compared to sham controls; elevated levels were decreased by incorporation of 2DG but not by a.a. in *Ccr2*^*RFP/+*^*Cx3cr1*^*GFP/+*^ mice. (Fig. 1c-f; Supplementary Fig. 1b). Among *Ccr2*^*RFP/RFP*^*Cx3cr1*^*GFP/GFP*^ mice (Supplementary Fig. 2b), there were no notable differences in CCR2- or CX3CR1-expression between sham (Supplementary Video 5) and aPLA groups (Supplementary Video 6), and flow cytometry data corroborated our visual observations (Supplementary Fig. 2c-d). Interestingly, CCR2^+^CX3CR1^+^ cells were decreased with aPLA implantation relative to sham controls in *Ccr2*^*RFP/RFP*^*Cx3cr1*^*GFP/GFP*^ mice (Supplementary Fig. 2e).

Next, we sought to elucidate the composition of the different CCR2^+^, CX3CR1^+^ and CCR2^+^CX3CR1^+^ populations in the implant microenvironment based on CD86 and CD206 expression. We classified proinflammatory populations^24^ as CD86^+^CD206^-^, anti-inflammatory populations^25^ as CD206^+^, and transition populations moving from proinflammatory to anti-inflammatory states^26,27^ as CD86^+^CD206^+^. Among CCR2^+^, CX3CR1^+^ and CCR2^+^CX3CR1^+^ populations, the proportion of proinflammatory cells were higher in the aPLA group compared to sham controls in *Ccr2*^*RFP/+*^*Cx3cr1*^*GFP/+*^ mice (Fig. 1g-i), whereas there were no differences with *Ccr2*^*RFP/RFP*^*Cx3cr1*^*GFP/GFP*^ mice (Supplementary Fig. 2f-h). Incorporating a.a. decreased proinflammatory levels among CX3CR1^+^ and CCR2^+^CX3CR1^+^ populations in *Ccr2*^*RFP/+*^*Cx3cr1*^*GFP/+*^ mice (Fig. 1h-i). The proportion of anti-inflammatory cells among CCR2^+^ cells was decreased with aPLA implantation compared to sham controls in *Ccr2*^*RFP/+*^*Cx3cr1*^*GFP/+*^ mice, while incorporating a.a. restored the proportion of these cells to a similar level as the sham controls (Fig. 1j).

However, no difference was observed in the proportion of anti-inflammatory CCR2^+^ cells in *Ccr2*^*RFP/RFP*^*Cx3cr1*^*GFP/GFP*^ mice (Supplementary Fig. 2i). Proportions of anti-inflammatory cells among CX3CR1^+^ and CCR2^+^CX3CR1^+^ populations were similar across all groups in *Ccr2*^*RFP/+*^*Cx3cr1*^*GFP/+*^ mice (Fig. 1k-l) and *Ccr2*^*RFP/RFP*^*Cx3cr1*^*GFP/GFP*^ mice (Supplementary Fig. 2j-k). Among, CCR2^+^ populations, the proportion of transition cells was reduced by the implantation of aPLA compared to sham controls in *Ccr2*^*RFP/+*^*Cx3cr1*^*GFP/+*^ mice (Fig. 1m), but not *Ccr2*^*RFP/RFP*^*Cx3cr1*^*GFP/GFP*^ mice (Supplementary Fig. 2l). Incorporation of either a.a. or 2DG increased proportions of transition cells (Fig. 1m). Similar to *Ccr2*^*RFP/+*^*Cx3cr1*^*GFP/+*^ mice (Fig. 1n-o), *Ccr2*^*RFP/RFP*^*Cx3cr1*^*GFP/GFP*^ mice (Supplementary Fig. 2m-n) showed no notable differences in proportions of transition cells among CX3CR1^+^ and CCR2^+^CX3CR1^+^ populations.

In comparison to sham controls, the proportion of CD45^+^ cells was increased by implantation of aPLA in *Ccr2*^*RFP/+*^*Cx3cr1*^*GFP/+*^ mice, while incorporating either a.a. or 2DG significantly diminished this effect (Fig. 1p). In contrast, the proportion of CD45^+^ cells decreased with aPLA implantation in *Ccr2*^*RFP/RFP*^*Cx3cr1*^*GFP/GFP*^ mice compared to their sham controls (Supplementary Fig. 2o). In both *Ccr2*^*RFP/+*^*Cx3cr1*^*GFP/+*^ mice and *Ccr2*^*RFP/RFP*^*Cx3cr1*^*GFP/GFP*^ mice, we observed increased fold change of proinflammatory CD45^+^ cells with respect to transition cells or anti-inflammatory cells in the aPLA group when compared to sham controls (Fig. 1q-r; Supplementary Fig. 2p-q). Also, the fold change of transition or anti-inflammatory CD45^+^ cells with respect to proinflammatory cells was decreased in the aPLA group when compared to sham controls (Fig. 1s-t; Supplementary Fig. 2r-s). Interestingly, incorporating a.a. had the dual effect of decreasing proinflammatory and increasing anti-inflammatory proportions of cells (Fig. 1q-t). In comparison to sham controls, Ly6G^+^ neutrophils^15^ were increased following aPLA implantation in *Ccr2*^*RFP/+*^*Cx3cr1*^*GFP/+*^ mice (Fig. 1u), whereas there were no changes in *Ccr2*^*RFP/RFP*^*Cx3cr1*^*GFP/GFP*^ mice (Supplementary Fig. 2t). Incorporating either a.a. or 2DG decreased elevated neutrophil levels (Fig. 1u).

### The composition of myeloid cells is reorganized toward pro-regenerative states by immunometabolic cues in the aPLA microenvironment

Having observed that CD45^+^ populations were strikingly different among groups in our study (Fig. 1p), we sought to uncover the relative contribution of myeloid cells, including CD11b^+^ monocytes^28-30^ (CD11b could also be expressed on subsets of B-cells, neutrophils and macrophages^31^), F4/80^+^ macrophages^31^ and CD11c^+^ dendritic cells^32,33^ to the constitution and polarization of the biomaterial microenvironment. In comparison to sham controls, we observed elevated monocyte populations following aPLA implantation in *Ccr2*^*RFP/+*^*Cx3cr1*^*GFP/+*^ mice, but not in *Ccr2*^*RFP/RFP*^*Cx3cr1*^*GFP/GFP*^ mice (Fig. 2a-b; Supplementary Fig. 2u). Incorporating either a.a. or 2DG reduced the frequency of monocytes compared to aPLA alone (Fig. 2a-b). Interestingly, there was increased fold change of proinflammatory cells with respect to transition cells or anti-inflammatory monocytes in the aPLA group when compared to sham controls, with either a.a. or 2DG incorporation modulating proinflammatory monocyte levels (Fig. 2c-d). Furthermore, the fold change of transition or anti-inflammatory monocytes with respect to proinflammatory cells was decreased in the aPLA group when compared to sham controls, with either a.a. or 2DG increasing anti-inflammatory monocyte levels (Fig. 2e-f).

**Figure 2.**
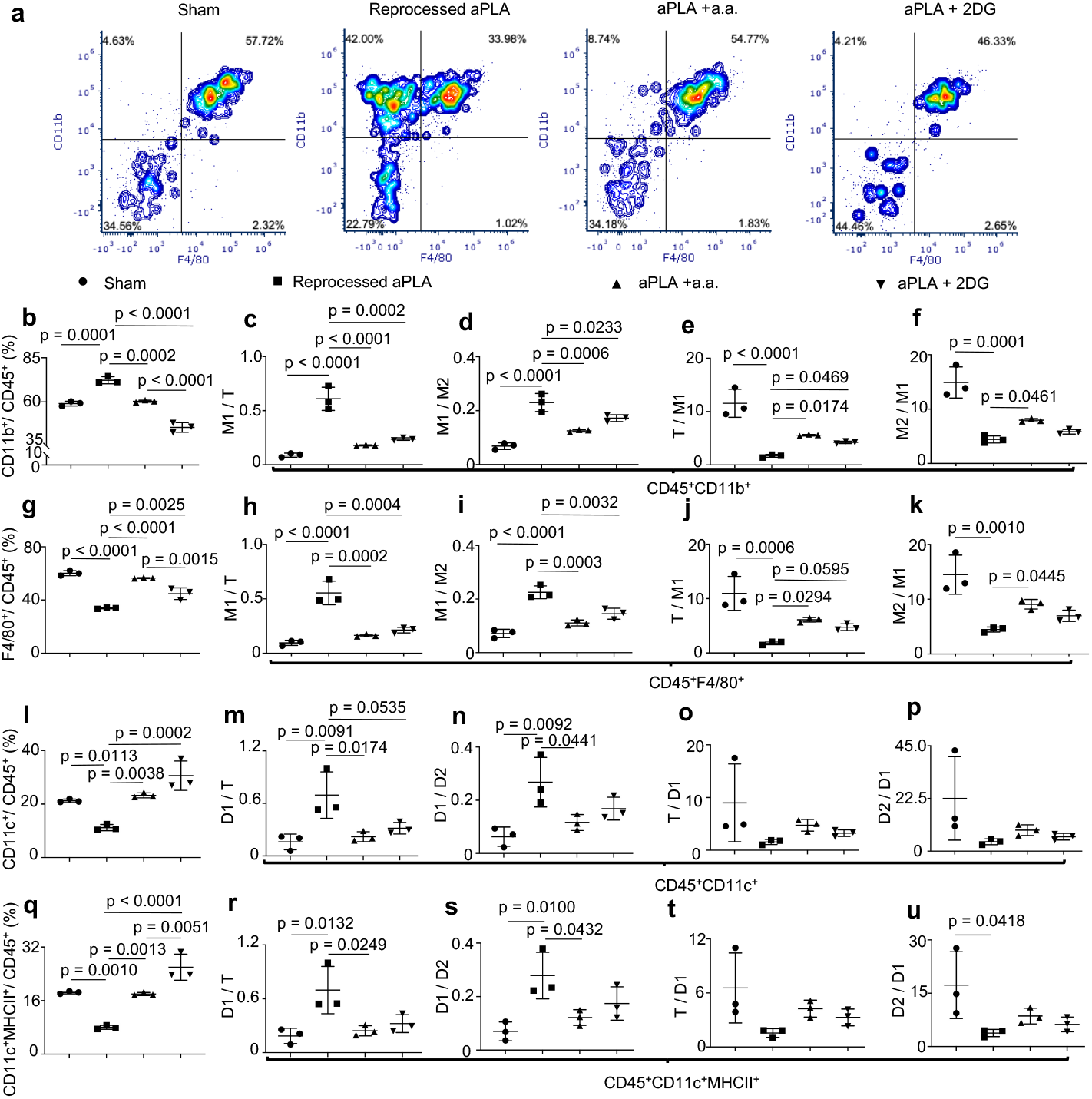
Polarization states of myeloid cells are regulated by targeting immunometabolism in the amorphous polylactide biomaterial microenvironment. **a**, Representative flow cytometry plots gated on CD45. **b**, Monocytes (CD45^+^CD11b^+^ cells). **c-d**, Fold change of proinflammatory (M1; CD86^+^CD206^-^) monocytes with respect to transition (T; CD86^+^CD206^+^) monocytes (c) or anti-inflammatory (M2; CD206^+^) monocytes (d). **e-f**, Fold change of T (e) or M2 (f) monocytes with respect to M1 monocytes. **g**, Macrophages (CD45^+^F4/80^+^ cells). **h-i**, Fold change of proinflammatory (M1; CD86^+^CD206^-^) macrophages with respect to transition (T; CD86^+^CD206^+^) macrophages (h) or anti-inflammatory (M2; CD206^+^) macrophages (i). **j-k**, Fold change of T (j) or M2 (k) macrophages with respect to M1 macrophages. **l**, Dendritic (CD45^+^CD11c^+^) cells. **m-n**, Fold change of proinflammatory (D1; CD86^+^CD206^-^) dendritic cells with respect to transition (T; CD86^+^CD206^+^) dendritic cells (m) or anti-inflammatory (D2; CD206^+^) dendritic cells (n). **o-p**, Fold change of T (o) or D2 (p) dendritic cells with respect to D1 dendritic cells. **q**, Dendritic cells expressing class II major histocompatibility complex (MHC II) molecules (CD45^+^CD11c^+^MHCII^+^ cells). **r-s**, Fold change of proinflammatory (D1; CD86^+^CD206^-^) MHCII^+^ dendritic cells with respect to transition (T; CD86^+^CD206^+^) MHCII^+^ dendritic cells (r) or anti-inflammatory (D2; CD206^+^) MHCII^+^ dendritic cells (s). **t-u**, Fold change of T (t) or D2 (u) MHCII^+^ dendritic cells with respect to D1 MHCII^+^ dendritic cells. One-way ANOVA followed by Tukey’s or Newman-Keul’s multiple comparison test, n = 3 mice per group; amorphous polylactide, aPLA; aminooxyacetic acid, a.a.; 2-deoxyglucose, 2DG.

We observed decreased macrophage expression in the aPLA group compared to sham controls of *Ccr2*^*RFP/+*^*Cx3cr1*^*GFP/+*^ or *Ccr2*^*RFP/RFP*^*Cx3cr1*^*GFP/GFP*^ mice, whereby incorporating either a.a. or 2DG increased macrophage expression (Fig. 2a, 2g; Supplementary Fig. 2v). Yet, exploring polarization states revealed increased fold change of proinflammatory macrophages with respect to transition or anti-inflammatory macrophages in the aPLA group when compared to sham controls, with either a.a. or 2DG incorporation reducing proinflammatory macrophage expression (Fig. 2h-i). Moreover, the fold change of transition or anti-inflammatory macrophages with respect to proinflammatory macrophages was decreased in the aPLA group when compared to sham controls, with a.a. increasing anti-inflammatory levels (Fig. 2j-k).

Dendritic cell populations were decreased in the aPLA group compared to sham controls of *Ccr2*^*RFP/+*^*Cx3cr1*^*GFP/+*^ mice, but increased in *Ccr2*^*RFP/RFP*^*Cx3cr1*^*GFP/GFP*^ mice (Fig. 2l; Supplementary Fig. 2w). Incorporating either a.a. or 2DG increased dendritic cell expression (Fig. 2l). Furthermore, we observed increased fold change of proinflammatory dendritic cells with respect to transition or anti-inflammatory dendritic cells in the aPLA group when compared to sham controls, with a.a. reducing proinflammatory expression (Fig. 2m-n). There were no differences in the fold change of transition or anti-inflammatory dendritic cells with respect to proinflammatory dendritic cells between groups (Fig. 2o-p). Observed trends in dendritic cells were similar to results of dendritic cells expressing class II major histocompatibility complex (MHCII) molecules (Fig. 2q-u; Supplementary Fig. 2x).

### Leveraging immunometabolism with aPLA favorably compares with currently used techniques

To determine how inhibiting the metabolism of immune cells using 2DG or a.a. compares to clinically used neutralization strategies^34^, we fabricated aPLA incorporating hydroxyapatite (HA)^20^. Wild-type B6 mice were surgically incised as sham controls or implanted with 1mm long aPLA formulations (Fig. 3a). Flow cytometric analyses demonstrated elevated neutrophils with aPLA compared to sham controls among CD45^+^ cells; elevated levels were decreased by either a.a. or 2DG, but not by HA (Fig. 3b). There was increased fold change of proinflammatory CD45^+^ cells with respect to transition cells or anti-inflammatory cells in the aPLA group when compared to sham controls (Fig. 3c-d). Incorporating a.a., 2DG or HA each decreased the fold change of proinflammatory CD45^+^ cells with respect to transition cells (Fig. 3c), but only a.a. or 2DG decreased the fold change of proinflammatory cells with respect to anti-inflammatory cells (Fig. 3d). The fold change of transition or anti-inflammatory CD45^+^ cells with respect to proinflammatory cells was decreased in the aPLA group when compared to sham controls (Fig. 3e-f). Whereas, incorporating a.a. or 2DG increased transition and anti-inflammatory expression, HA did not (Fig. 3e-f). Importantly, a.a. had greater effects than HA at decreasing proinflammatory and increasing transition or anti-inflammatory CD45^+^ proportions (Fig. 3c-f).

**Figure 3.**
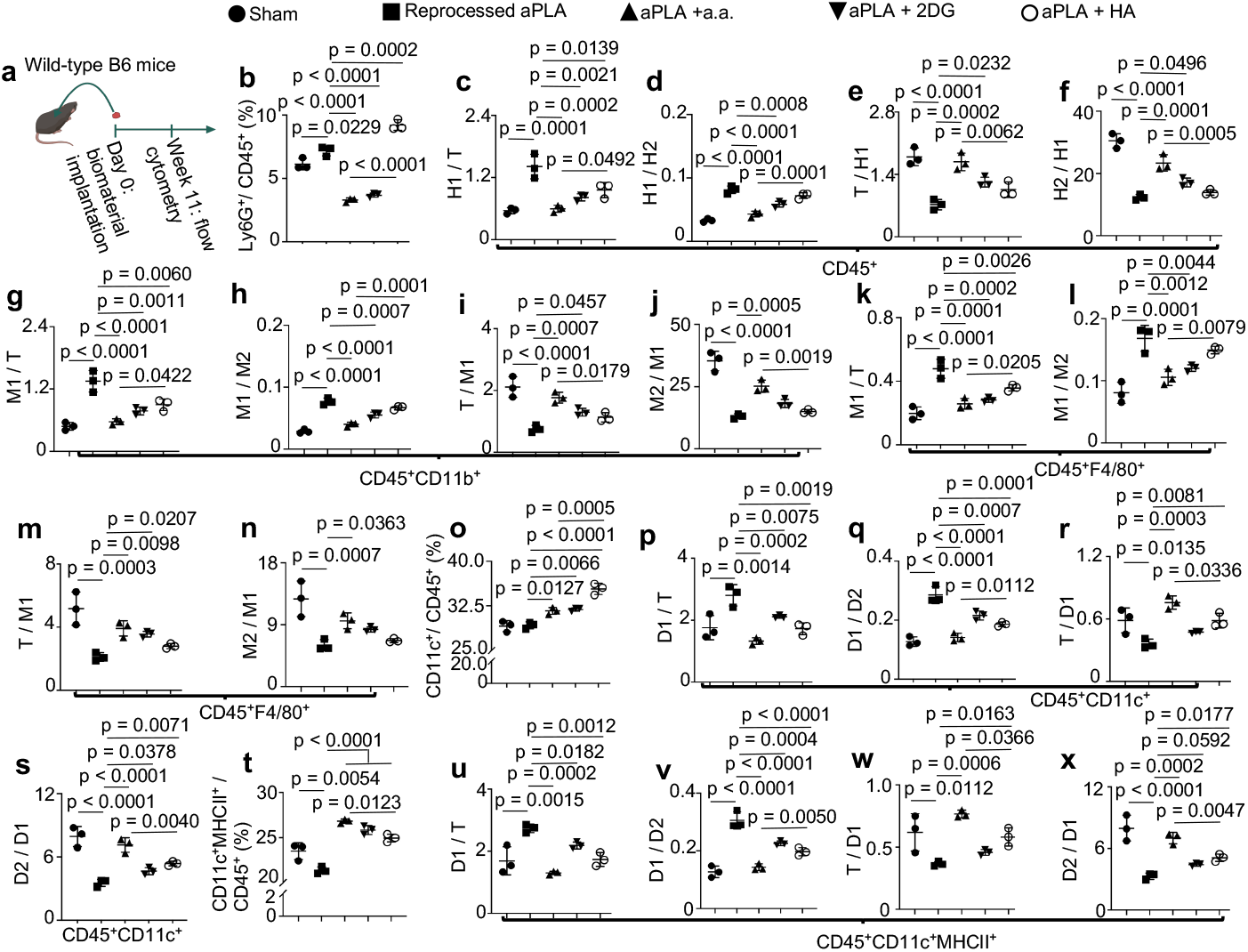
Using an acid more favorably modulates activation states of immune cells around amorphous polylactide biomaterials compared to traditional neutralization techniques. **a**, Wild-type B6 mice were surgically incised (sham group) or implanted with reprocessed amorphous polylactide (aPLA), aPLA incorporating aminooxyacetic acid (a.a.), 2-deoxyglucose (2DG) or hydroxyapatite (HA). Afterwards, flow cytometric analysis of tissues around incision sites (sham group) or biomaterials was undertaken. **b**, Neutrophils (CD45^+^Ly6G^+^ cells). **c-d**, Fold change of proinflammatory (H1; CD86^+^CD206^-^) cells with respect to transition (T; CD86^+^CD206^+^) cells (c) or anti-inflammatory (H2; CD206^+^) cells (d), gated for nucleated hematopoietic (CD45^+^) populations. **e-f**, Fold change of T (e) or H2 (f) cells with respect to H1 cells. **g-h**, Fold change of proinflammatory (M1; CD86^+^CD206^-^) monocytes (CD45^+^CD11b^+^) with respect to transition (T; CD86^+^CD206^+^) monocytes (g) or anti-inflammatory (M2; CD206^+^) monocytes (h). **i-j**, Fold change of T (i) or M2 (j) monocytes with respect to M1 monocytes. **k-l**, Fold change of proinflammatory (M1; CD86^+^CD206^-^) macrophages (CD45^+^F4/80^+^) with respect to transition (T; CD86^+^CD206^+^) macrophages (k) or anti-inflammatory (M2; CD206^+^) macrophages (l). **m-n**, Fold change of T (m) or M2 (n) macrophages with respect to M1 macrophages. **o**, Dendritic (CD45^+^CD11c^+^) cells. **p-q**, Fold change of proinflammatory (D1; CD86^+^CD206^-^) dendritic cells with respect to transition (T; CD86^+^CD206^+^) dendritic cells (p) or anti-inflammatory (D2; CD206^+^) dendritic cells (q). **r-s**, Fold change of T (r) or D2 (s) dendritic cells with respect to D1 dendritic cells. **t**, Dendritic cells expressing class II major histocompatibility complex (MHC II) molecules (CD45^+^CD11c^+^MHCII^+^ cells). **u-v**, Fold change of proinflammatory (D1; CD86^+^CD206^-^) MHCII^+^ dendritic cells with respect to transition (T; CD86^+^CD206^+^) MHCII^+^ dendritic cells (u) or anti-inflammatory (D2; CD206^+^) MHCII^+^ dendritic cells (v). **w-x**, Fold change of T (w) or D2 (x) MHCII^+^ dendritic cells with respect to D1 MHCII^+^ dendritic cells. One-way ANOVA followed by Tukey’s multiple comparison test, n = 3 mice per group.

The fold change of proinflammatory monocytes with respect to transition or anti-inflammatory monocytes was increased in the aPLA group when compared to sham controls (Fig. 3g-h). Elevated proinflammatory monocytes proportions were consistently decreased by incorporating a.a. or 2DG, with HA only decreasing elevated proportions with respect to transition cells (Fig. 3g-h). On the other hand, the fold change of transition or anti-inflammatory monocytes with respect to proinflammatory monocytes was decreased in the aPLA group when compared to sham controls (Fig. 3i-j). Whereas incorporating a.a. or 2DG increased the proportion of transition monocytes, only a.a. increased the proportion of anti-inflammatory monocytes (Fig. 3i-j). Notably, a.a. was more effective than HA at consistently reducing proinflammatory and increasing transition or anti-inflammatory monocyte proportions (Fig. 3g-i).

With macrophages, the fold change of proinflammatory cells with respect to transition or anti-inflammatory cells was increased in the aPLA group when compared to sham controls (Fig. 3k-l). Increased proinflammatory macrophage proportions were reduced by incorporating a.a. or 2DG, with HA only decreasing elevated proportions with respect to transition cells (Fig. 3k-l). We observed that a.a. was more effective than HA at reducing proinflammatory macrophage proportions (Fig. 3k-l). Additionally, the fold change of transition or anti-inflammatory macrophages with respect to proinflammatory macrophages was decreased in the aPLA group when compared to sham controls (Fig. 3m-n). As with monocytes, incorporating a.a. or 2DG increased the proportion of transition macrophages, with only a.a. increasing the proportion of anti-inflammatory macrophages (Fig. 3m-n).

While aPLA implantation did not reduce dendritic cell populations when compared to sham controls, incorporating a.a., 2DG or HA increased dendritic cell populations (Fig. 3o). Interestingly, the fold change of proinflammatory dendritic cells with respect to transition or anti-inflammatory dendritic cells was increased in the aPLA group when compared to sham controls, and incorporating a.a., 2DG or HA decreased elevated proinflammatory proportions (Fig. 3p-q). We also observed that the fold change of transition or anti-inflammatory dendritic cells with respect to proinflammatory dendritic cells was decreased in the aPLA group when compared to sham controls (Fig. 3r-s). Incorporating a.a. or HA increased the proportion of transition dendritic cells, with a.a., 2DG or HA increasing the proportion of anti-inflammatory dendritic cells (Fig. 3r-s). When compared with HA, a.a. decreased proinflammatory, and increased transition or anti-inflammatory proportions (Fig. 3q-s).

The expression of MHCII^+^ dendritic cells was decreased with aPLA implantation when compared to sham controls, with incorporating a.a., 2DG or HA increasing MHCII^+^ dendritic cell expression (Fig. 3t). The fold change of proinflammatory MHCII^+^ dendritic cells with respect to transition or anti-inflammatory MHCII^+^ dendritic cells was increased in the aPLA group when compared to sham controls, and incorporating a.a., 2DG or HA decreased elevated proinflammatory proportions (Fig. 3u-v). Also, the fold change of transition or anti-inflammatory MHCII^+^ dendritic cells with respect to proinflammatory MHCII^+^ dendritic cells was decreased in the aPLA group when compared to sham controls (Fig. 3w-x). Incorporating a.a. or HA increased the proportion of transition MHCII^+^ dendritic cells, with a.a., 2DG or HA increasing the proportion of anti-inflammatory MHCII^+^ dendritic cells (Fig. 3w-x). In comparison to HA, a.a. decreased proinflammatory, and increased transition or anti-inflammatory proportions (Fig. 3v-x).

### Immunometabolic cues from crystalline polylactide create a pro-regenerative microenvironment with the help of T-cells

In comparison to aPLA, we have observed by electrospray ionization-mass spectrometry that (semi-) crystalline polylactide (cPLA) formulations degrade more slowly^20^. Furthermore, the different stereochemical compositions of aPLA and cPLA could elicit differential immune cellular responses by triggering varied bioenergetic signatures^7^. As such, we sought to uncover the composition and phenotypes of immune cells in the cPLA microenvironment. Wild-type B6 mice were surgically incised as sham controls; implanted with 1mm long cPLA formulations with and without incorporating a.a. and 3-(3-pyridinyl)-1-(4-pyridinyl)-2-propen-1-one (3PO; Fig 4a), with 3PO being a small molecule inhibitor of 6-phosphofructo-2-kinase^35^, the rate limiting enzyme in glycolysis. Flow cytometric analyses demonstrated that there was increased CD45 expression with cPLA implantation in comparison to sham controls, and elevated levels were decreased by incorporating a.a. (Fig. 4b). There was increased fold change of proinflammatory CD45^+^ cells with respect to transition cells or anti-inflammatory cells in the cPLA group when compared to sham controls (Supplementary Fig. 3a-b). Also, the fold change of transition or anti-inflammatory CD45^+^ cells with respect to proinflammatory cells was decreased in the cPLA group when compared to sham controls (Supplementary Fig. 3c-d). Interestingly, incorporation of a.a. or 3PO did not decrease proinflammatory nor increase transition or anti-inflammatory CD45^+^ proportions (Supplementary Fig. 3a-d).

**Figure 4.**
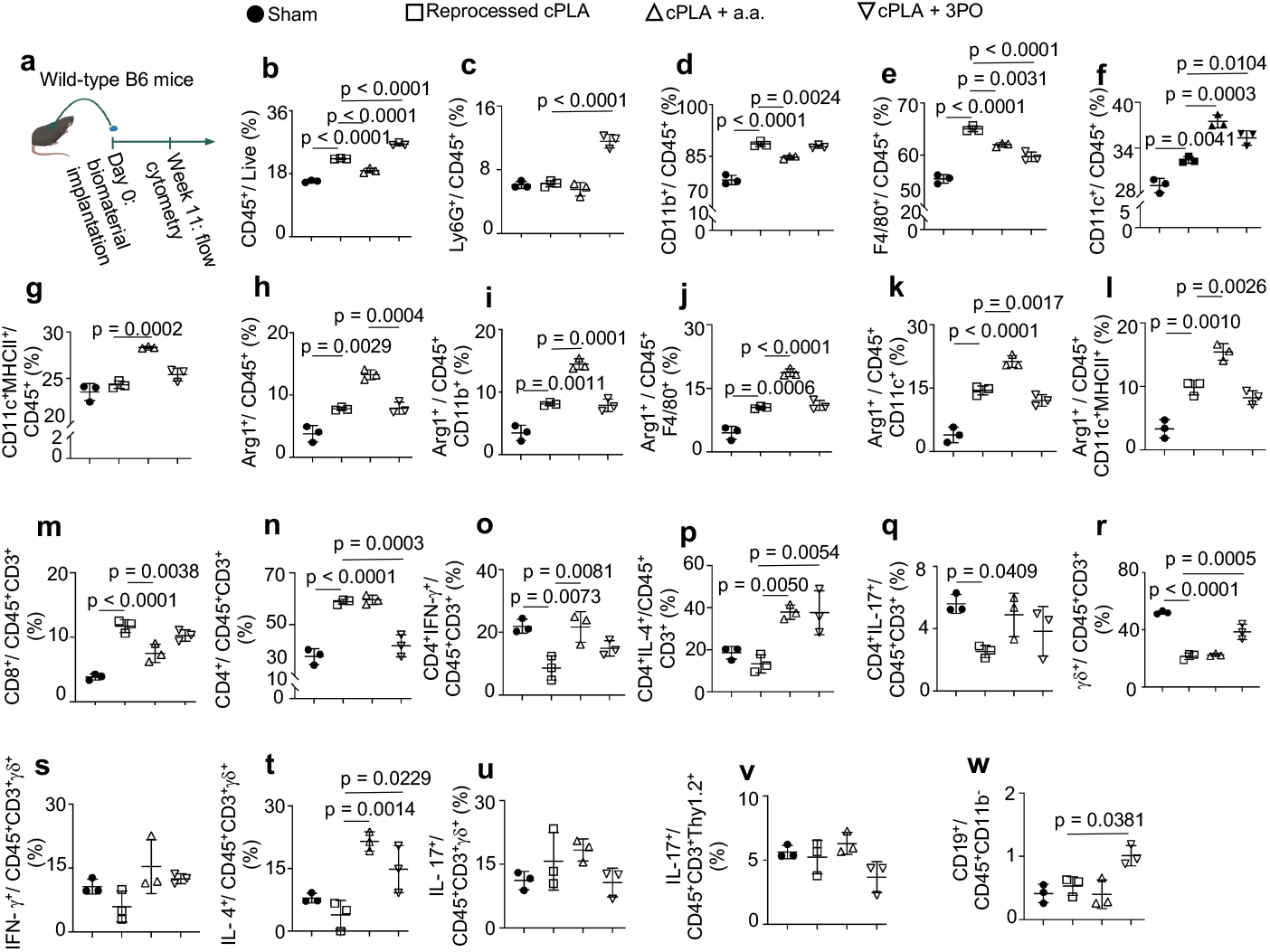
Locally targeting immunometabolism in the crystalline polylactide environment elevates interleukin-4 (IL-4)-expressing T cell subsets with differential effects on myeloid populations. **a**, Wild-type B6 mice were surgically incised (sham group) or implanted with reprocessed crystalline polylactide (cPLA), cPLA incorporating aminooxyacetic acid (a.a.) or 3-(3-pyridinyl)-1-(4-pyridinyl)-2-propen-1-one (3PO). Afterwards, flow cytometric analysis of tissues around incision sites (sham group) or biomaterials was undertaken. **b**, Nucleated hematopoietic (CD45^+^) cells. **c**, Neutrophils (CD45^+^Ly6G^+^ cells). **d**, Monocytes (CD45^+^CD11b^+^ cells). **e**, Macrophages (CD45^+^F4/80^+^ cells). **f**, Dendritic (CD45^+^CD11c^+^) cells. **g**, Dendritic cells expressing class II major histocompatibility complex (MHC II) molecules (CD45^+^CD11c^+^MHCII^+^ cells). **h-l**, Nucleated hematopoietic cells (h), monocytes (i), macrophages (j), dendritic cells (k), MHCII^+^ dendritic cells (l) expressing Arginase 1 (Arg1^+^). **m**, Cytotoxic T lymphocytes (CD45^+^CD3^+^CD8^+^ cells). **n**, T helper lymphocytes (CD45^+^CD3^+^CD4^+^ cells). **o**, T helper 1 cells expressing interferon-gamma (IFN-γ). **p**, T helper 2 cells expressing interleukin-4. **q**, T helper 17 cells expressing interleukin-17A. **r**, gamma delta (γδ) T (CD45^+^CD3^+^γδ+) cells. **s-u**, γδ+ T cells producing IFN-γ (s), IL-4 (t) and IL-17A (u). **v**, Innate lymphoid cells (CD45^+^CD3^+^Thy1.2^+^) producing IL-17A. **w**, B cells (CD45^+^CD11b^-^CD19^+^). One-way ANOVA followed by Tukey’s multiple comparison test, n = 3 mice per group.

There was no difference in neutrophil expression in the cPLA group when compared with sham controls (Fig. 4c). We observed increased monocyte expression in the cPLA group when compared to sham controls; elevated monocyte expression was decreased by incorporating a.a. (Fig. 4d). While there was increased fold change of proinflammatory monocytes with respect to transition or anti-inflammatory monocytes, incorporation of a.a. or 3PO did not reduce elevated proinflammatory levels (Supplementary Fig. 3e-f). The fold change of transition or anti-inflammatory monocytes with respect to proinflammatory monocytes was decreased in the cPLA group when compared to sham controls, and incorporation of a.a. or 3PO did not have significant effects (Supplementary Fig. 3g-h).

Macrophage expression was increased by cPLA implantation in comparison to sham controls, with incorporation of a.a. or 3PO reducing elevated expression (Fig. 3e). Furthermore, there was increased fold change of proinflammatory macrophages with respect to transition or anti-inflammatory macrophages in the cPLA group when compared to sham controls (Supplementary Fig. 3i-j). The fold change of transition or anti-inflammatory macrophages with respect to proinflammatory macrophages was decreased in the cPLA group when compared to sham controls (Supplementary Fig. 3k-l). Incorporation of a.a. or 3PO neither decreased proinflammatory nor increased anti-inflammatory proportions (Supplementary Fig. 3i-l).

Dendritic cell levels were increased in the cPLA group in comparison to sham controls, with a.a. or 3PO further increasing dendritic cell expression (Fig. 4f). Whereas there were no changes in the fold change of proinflammatory dendritic cells with respect to transition cells, the fold change of proinflammatory dendritic cells with respect to anti-inflammatory cells was increased in the cPLA group when compared to sham controls, with incorporation of 3PO reducing elevated levels (Supplementary Fig. 3m-n). The fold change of transition or anti-inflammatory dendritic cells with respect to proinflammatory cells was decreased in the cPLA group when compared to sham controls (Supplementary Fig. 3o-p).

The expression of MHCII^+^ dendritic cells with cPLA implantation was similar to sham controls, and MHCII^+^ expression was increased by incorporation of a.a. (Fig. 4g). In addition, the fold change of proinflammatory MHCII^+^ dendritic cells with respect to transition or anti-inflammatory MHCII^+^ dendritic cells was increased in the cPLA group when compared to sham controls (Supplementary Fig. 3q-r). Also, the fold change of transition or anti-inflammatory MHCII^+^ dendritic cells with respect to proinflammatory MHCII^+^ dendritic cells was decreased in the cPLA group when compared to sham controls (Supplementary Fig. 3s-t). Incorporation of a.a. or 3PO neither decreased proinflammatory nor increased anti-inflammatory levels (Supplementary Fig. q-t). While Arginase 1 (Arg1) levels were increased in the cPLA group when compared to sham controls, incorporation of a.a. further increased Arg 1 levels among CD45^+^, CD11b^+^, F4/80^+^, CD11c^+^ and CD11c^+^MHCII^+^ populations (Fig. 4h-l).

Given the distinct observation made with cPLA implants, we sought to elucidate the role of the adaptive immune response, including CD19 B cell activity^36^, CD4 T helper and CD8 cytotoxic T cells^25,37^. With cPLA implantation, CD8 expression was increased when compared to sham controls, and incorporation of a.a. decreased elevated levels (Fig. 4m). Similarly, CD4 expression was higher in the cPLA group compared to sham controls, with 3PO reducing elevated levels (Fig. 4n). CD4^+^ cells expressing IFN-γ were reduced in the cPLA group when compared to sham controls; incorporation of a.a. or 3PO tended to increase IFN-γ expression only to levels similar to the sham group (Fig. 4o). In contrast, CD4^+^ cells expressing IL-4 were similar between the cPLA group and sham controls, with incorporation of a.a. or 3PO increasing IL-4 levels (Fig. 4p). Assessing CD4^+^ cells expressing IL-17 revealed reduced expression in the cPLA group when compared to sham controls; incorporation of a.a. or 3PO tended to increase IL-17 expression only to levels similar to the sham group (Fig. 4q). While the γδ-repertoires was decreased in the cPLA group when compared to sham controls, incorporation of 3PO but not a.a. tended to increase the T cell receptor repertoires (Fig. 4r). Additionally, while there were no changes in IFN-γ and IL-17 expression from γδ^+^ T-cells, incorporation of a.a. or 3PO increased IL-4 expression from γδ^+^ T-cells (Fig. 4s-u). There were no changes in IL-17 expression from innate lymphoid cells (Fig. 4v). We did not observe changes in CD19 expression between the cPLA group and sham controls; however, incorporation of 3PO tended to increase CD19 expression (Fig. 4w).

## Discussion

Although polylactide-based medical devices are FDA-approved for applications in adult reconstructive surgery, drug delivery and nanotechnology, their clinical utility is significantly limited by long-term, sterile inflammation, which is poorly understood^3,38^. Our findings suggest that the activation states and trafficking of immune cells to the amorphous polylactide (aPLA) biomaterial microenvironment is dependent on CCR2 and CX3CR1 expression, with CCR2^5^ likely to play a greater role over CX3CR1^18,19^ expression. We reveal that both CCR2 and CX3CR1 signaling could be regulated by locally controlling glycolytic flux through targeting hexokinase in the biomaterial microenvironment. In liver models of sterile injury as well as in the heart following myocardial infarction, CCR2^+^ monocytes dominate the proinflammatory phase, with CCR2^+^CX3CR1^+^ and CX3CR1^+^ monocytes and macrophages playing a greater role during the anti-inflammatory or reparative phase of healing^14,39^. In our study, however, sterile aPLA implants increase proinflammatory (CD86^+^CD206^-^) proportions of CCR2^+^, CX3CR1^+^ and CCR2^+^CX3CR1^+^ cell populations, while decreasing transition (CD86^+^CD206^+^) and anti-inflammatory proportions (CD206^+^) of CCR2^+^ populations, an effect that required CCR2- and CX3CR1-competency. Against our hypothesis, incorporation of a.a did not reduce monocyte recruitment around biomaterials; however, it reshaped the composition of recruited CCR2^+^, CX3CR1^+^ and CCR2^+^CX3CR1^+^ cell populations to comprise reduced proinflammatory, and elevated transition and anti-inflammatory proportions. These transition immune cell populations suggest a reversal of proinflammatory biomaterial responses, playing a crucial role in angiogenesis and tissue regeneration^26,27^. Consistent with our findings, an increase in CX3CR1^+^ populations is observed around polylactide copolymers^40^.

Myeloid cell recruitment to aPLA-based implants appears to be regulated by immunometabolism in the biomaterial microenvironment. We observed a two-pronged effect of aPLA in constituting a proinflammatory biomaterial microenvironment—aPLA elevated the relative levels of proinflammatory monocytes while concurrently decreasing the relative proportions of transition and anti-inflammatory monocytes. This trend is observable in-vitro with polyethylene wear particles^6,10^ and other types of polylactide materials^7^, whose implantation result in chronic inflammatory responses. Consequently, it is likely that this two-pronged effect occurs with other classes of biomaterials, providing new insight into complex ways that biomaterials modify their immune microenvironment.

During the inflammatory response to implants, the biomaterial microenvironment is glycolytically reprogrammed, showing enhanced radiolabeled glucose uptake in mice^20^, an observation made in implanted human patients^41,42^. Our findings reveal that disrupting elevated glycolytic flux using 2DG or a.a. decreased proinflammatory monocytes, increased transition and anti-inflammatory monocytes, and decreased neutrophil recruitment to the biomaterial microenvironment. Paradoxically, aPLA implantation decreased macrophage and dendritic cell recruitment, while lowering MHCII expression on dendritic cells, an observation that is thought to imply an immunomodulatory role of polylactide implants^32^. We reproduce these findings, but also uncover that the relative proportion of proinflammatory macrophages, dendritic cells or dendritic cells expressing MHCII is increased by aPLA implantation. With macrophages and, to lesser extent, dendritic cells and dendritic cells expressing MHCII, inhibiting glycolysis reduced proinflammatory and increased transition and anti-inflammatory proportions, providing new insight on how biomaterials could result in chronic inflammation all the while orchestrating seemingly conflicting immunological events. Of note, CCR2- and CX3CR1-deficiency prevents the recruitment of neutrophils, monocytes, macrophages, dendritic cells and MHCII expression on dendritic cells in the aPLA microenvironment, which may be due to the role of CCR2 in promoting the local activation and maturation of immune cells^43,44^.

Historically, chronic inflammation by aPLA is thought to be due to acidic degradation products^34^. As such, neutralizing aPLA products with alkaline salts, like hydroxyapatite (HA), is currently the mainstay in clinics. Contrary to HA’s effects, a.a. further makes the biomaterial microenvironment acidic^20^; therefore, in principle, a.a. should worsen aPLA-induced inflammation. We show that, while incorporating a.a. reduced neutrophil levels, HA increased neutrophil recruitment, accentuating inflammation. To extents greater than HA’s, incorporating a.a. in aPLA implants reduced the relative proportion of proinflammatory immune cells, including overall nucleated hematopoietic cell populations, monocytes, macrophages, dendritic cells and dendritic cells expressing MHCII. Furthermore, a.a. largely increased transition and anti-inflammatory cellular proportions more than HA could. This suggests that, while pH may exert some role in the process, it is not the sole regulator of immunological events in the polylactide microenvironment.

Crystalline polylactide (cPLA) degrades at significantly slower rates than aPLA^20^. In contrast to observations made with aPLA, cPLA increased macrophage and dendritic cell recruitment, did not elevate neutrophil levels nor reduce MHCII expression on recruited dendritic cells. Also, unlike with aPLA, glycolytic inhibition did not reduce myeloid proinflammatory nor increase myeloid transition and anti-inflammatory proportions. However, glycolytic inhibition in the cPLA microenvironment reduced macrophage recruitment, specifically increased IL-4-expressing γδ^+^ T-cells and T helper 2 cells, while keeping IFN-γ at homeostatic levels, necessary for tissue physiology^45^. Importantly, a.a. reduced monocyte recruitment and increased Arg1 expression among overall myeloid populations, including monocytes, macrophages, dendritic cells and dendritic cells expressing MHCII. This observation is consistent with a.a. inhibiting aspartate aminotransferase, a key transaminase in the aspartate-arginosuccinate shunt in proinflammatory macrophages during metabolic reprogramming^46^. Antigen presenting cells are able to activate both class I and II MHC, following exposure to biomaterials^47^. While a.a. reduced cytotoxic T lymphocytes, 3PO reduced overall T helper cell recruitment to the cPLA biomaterial microenvironment, consistent with the crucial role of metabolism on T-cell function^48^.

In summary, our findings provide new insight on the role of immunometabolic cues on immune cellular trafficking to the biomaterial microenvironment, and how this affects the composition and activation states of immune cell populations. As orally-delivered small molecules, both a.a. and 3PO, as well as their derivatives, have been well tolerated when administered during clinical trials in cancer and Huntington’s disease^23,49^. Therefore, temporally-regulated, local release of these metabolic modulators from implants to program the trafficking and polarization of immune cells in the biomaterial microenvironment offers a highly translatable opportunity that could advance regenerative engineering toward improved human and animal health.

## Methods

### Biomaterial formulation and metabolic modulators

Amorphous polylactide (aPLA; PLA 4060D) and semi-crystalline polylactide (cPLA; PLA 3100HP) biomaterials from NatureWorks LLC were used. As metabolic modulators and for neutralization studies, 3-(3-pyridinyl)-1-(4-pyridinyl)-2-propen-1-one (3PO; MilliporeSigma), 2-deoxyglucose (2DG; MilliporeSigma), aminooxyacetic acid (a.a.; Sigma-Aldrich) and hydroxyapatite (HA; 2.5 μm^2^ particle sizes^50^; Sigma-Aldrich) were incorporated into biomaterials by melt-blending at 190 ºC for 3 mins in a DSM 15 cc mini-extruder (DSM Xplore), then made into pellets using a pelletizer (Leistritz Extrusion Technology). Afterwards, pellets were made into 1.75 mm (diameter) filaments using an extruder (Filabot EX2) at 170 ºC with air set at 93. Filaments were cut into 1 mm-long or 7.5 mm-long sizes, then sterilized by ultraviolet radiation for 30 minutes^51^. Based on prior studies^20^, we estimated that 189 mg of 2DG, 4.86 mg of 3PO, 90 mg of a.a. or 200 mg of HA in 10 g of aPLA or cPLA will approximate concentrations that were effective when applied in-vitro. To control for melt-blending as a confounder, biomaterials not incorporating metabolic modulators or HA, were processed under similar conditions to make “reprocessed” formulations.

### Mice

Animal studies were approved by the Institutional Animal Care and Use Committee at Michigan State University (PROTO202100327). *Ccr2*^*RFP/RFP*^*Cx3cr1*^*GFP/GFP*^ mice, B6(Cg)-Tyr^c-2J^/J (B6 albino) mice and C57BL/6J (wild-type B6) mice were obtained from the Jackson Laboratory. To generate *Ccr2*^*RFP/+*^*Cx3cr1*^*GFP/+*^ mice, we crossed female *Ccr2*^*RFP/RFP*^*Cx3cr1*^*GFP/GFP*^ (8-week old) mice to male B6(Cg)-Tyr^c-2J^/J (B6 albino; 8-week old) mice, as previously described^18,19^. At 4-week old, generated *Ccr2*^*RFP/+*^*Cx3cr1*^*GFP/+*^ mice were assigned n = 3 (two females, one male) per group. Only female mice (n = 3 per group) were used for studies involving *Ccr2*^*RFP/RFP*^*Cx3cr1*^*GFP/GFP*^ mice (14-week old) and C57BL/6J mice (9-week old).

### Subcutaneous surgical model

Anesthesia was accomplished using isoflurane (2-3 %). Using aseptic technique, the skin of each mouse was shaved and disinfected using iodine and alcohol swabs. Surgical incision was made through the skin into the subcutis, with or without biomaterial implantation after a pouch had been made with forceps. Surgical glue (3M Vetbond) was used to close the skin, and each mouse received intraperitoneal or subcutaneous pre- and post-operative meloxicam (5 mg/ kg) injections as well as postoperative saline. In *Ccr2*^*RFP/+*^*Cx3cr1*^*GFP/+*^ mice and *Ccr2*^*RFP/RFP*^*Cx3cr1*^*GFP/GFP*^ mice, the neck (just caudal to the ear; Supplementary Fig. 1a) was surgically incised (sham) and implanted using 7.5 mm long filaments to allow for imaging of the biomaterial microenvironment. In C57BL/6J mice, the dorsum (back) of mice incised (sham) and implanted using 1 mm long filaments.

### Intravital microscopic imaging and processing

Mice were anesthetized using a isoflurane (2-3 %), and image stacks were acquired using a Leica SP8 DIVE laser multiphoton microscope equipped with Spectra-Physics Insight X3 dual beam (630 to 1300 nm tunable and 1040 nm fixed) and 4Tune, tunable, super sensitive hybrid detectors (HyDs). To acquire serial optical sections, a laser beam (940 nm for GFP; 1040 nm for RFP) was focused through a 25x water-immersion lens (NA 1.00 HC PL IRAPO, Leica) and scanned with a field of view of 0.59 × 0.59 mm^2^ at 600 Hz. To visualize a larger area, 3 tiles of optical fields were imaged using a motorized stage to automatically acquire sequential fields of view. Z-stacks were acquired in 3 μm steps to image a total depth of 117 μm of tissue. To avoid fluorophore bleed-through, images were acquired using sequential scanning in between frames. Visualization of collagen was achieved via the second harmonic signal using the blue channel at 940 nm. Raw image stacks were imported into Fiji software (v1.53t; National Institute of Health) for tile merging. The tiled images were stitched by a grid/collection stitching plugin in Fiji. The merged image stacks were then imported into Imaris software (v10.0.0; Bitplane/Oxford Instruments) for further processing. Melanin autofluorescence from mouse skin was subtracted from the green and red channels. Also, GFP-expressing cells in the epidermis were excluded from images, as these are likely dendritic epidermal T cells^52,53^. The filtered red, green, and blue channels were then z-projected and shown as a single 2D image, with videos created in Imaris to show the individual slices from processed z-stacks.

### Tissue harvesting and dissociation

After 11 weeks post-operatively, mice were shaved around incision sites (sham) or biomaterials, then euthanized to obtain biopsies. As some implants had migrated and were unidentifiable, biopsies were obtained from only visible implants. In *Ccr2*^*RFP/+*^*Cx3cr1*^*GFP/+*^ mice and *Ccr2*^*RFP/RFP*^*Cx3cr1*^*GFP/GFP*^ mice, rectangular biopsies (9.5 mm long and 3.75 mm wide) around incision sites (sham) or biomaterials were collected. In C57BL/6J mice, circular biopsies (8 mm diameter) were collected. Tissues from different mice belonging to the same group in each study were collected together for dissociation. Tissues were placed into 10 mL of an enzyme cocktail containing 0.5 mg/ mL Liberase (Sigma-Aldrich), 0.5 mg/ mL Collagenase Type IV (Stem Cell Technologies), 250 U/ mL Deoxyribonuclease I (Worthington Biochemical Corporation) in 25mM HEPES buffer (Sigma-Aldrich) on a serrated Petri dish. Next, tissues were cut with surgical scissors for ∼1 minute and moved to an incubator at 37°C with 5% CO2 on top of an orbital shaker, shaking at 70 rpm for 1 hour. After the incubation period, the petri dish was removed and 5 mL of the enzyme cocktail and dissociated cells were put through a 70 μm filter into a 50mL conical tube. In another 5 mL of enzyme cocktail, undigested (residual) tissues were mechanically dissociated by being pressed against the serrated portion of the petri dish. Afterwards, using a 25 mL pipette, the 5mL of enzyme cocktail was filtered into a 50mL conical. Any undigested tissue on top of the 70μm filter was further mechanically digested with the thumb press of a syringe plunger for optimal extraction of cells. Using the same 25mL pipette, 30mL of cold Hanks’ Balanced Salt Solution without calcium, magnesium and phenol red (ThermoFisher Scientific) was used to wash the digestion petri dish and filtered into the 50mL conical. The cells in the 50mL conical were centrifuged at 350 x g for 10 minutes and the supernatant was discarded. Sedimented cells were counted then used for flow cytometry.

### Flow cytometry

Following tissue digestion, for experiments involving *Ccr2*^*RFP/+*^*Cx3cr1*^*GFP/+*^ mice and *Ccr2*^*RFP/RFP*^*Cx3cr1*^*GFP/GFP*^ mice, 1.5x10^6^ cells /well (n = 3 wells) were used for staining in a polypropylene 96-well round bottom plate. All staining steps were performed in 100 μL volume in the dark at 4 °C. Samples were first incubated with LIVE/DEAD Fixable Blue Dead Cell Stain Kit (1:500, Thermofisher, L23105) for 30 min. Cells were washed once with flow buffer (1X phosphate buffered saline (PBS), 0.5% bovine serum albumin), followed by incubation with TruStain FcX PLUS (anti-mouse CD16/32) Antibody (BioLegend, 156603; 0.25 μg/sample) for 10 minutes. The following antibodies were mixed and added directly to the cell suspension: BV421 CD86 (1:200, Biolegend, 105031), PacBlue Ly6G (1:150, Biolegend, 127611), BV605 CD45 (1:300, Biolegend, 103139), BV785 F4/80 (1:300, Biolegend, 123141), PerCP MHCII (1:200, Biolegend, 107623), PE-Dazzle 594 CD11c (1:500, Biolegend, 117347), APC CD206 (1:200, Biolegend, 141707) and AF700 CD11b (1:400, Biolegend, 101222). Cells and antibody mixture were incubated for 30 minutes. Cells were washed twice with flow staining buffer and fixed with 4% PFA for 10 minutes and resuspended in a final volume of 100 μL for flow cytometry analysis.

For experiments involving C57BL/6J mice, 1x10^6^ cells were used for staining in a polypropylene 96-well round bottom plate (n = 3). All staining steps were performed in 100 μL volume in the dark at 4°C. Samples were first incubated with LIVE/DEAD Fixable Blue Dead Cell Stain Kit (1:500, Thermofisher, cat#L23105) for 20 minutes. Cells were washed once with flow buffer, followed by incubation with TruStain FcX (anti-mouse CD16/32) Antibody (BioLegend, 101319; 1 μg/sample) in 50 μL volume for 10 minutes. The following antibodies were mixed together and added directly to the cell suspension: BV605 CD45 (1:500, Biolegend, 103139), AF700 CD11b (1:300, Biolegend, 101222), BV785 F4/80 (1:300, Biolegend, 123141), BV421 CD86 (1:200, Biolegend, 105031), APC CD206 (1:200, Biolegend, 141707), PerCP MHCII (1:200, Biolegend, 107623), SparkBlue 550 CD3 (1:100, Biolegend, 100259), APC-Fire 810 CD4 (1:100, Biolegend, 100479), BB700 CD8a (1:100, Biolegend, 566410), BV711 γδ TCR (1:200, BD Bioscience, 563994), BV480 Thy 1.2 (CD90.2; 1:40, BD Bioscience, 746840), BUV615 CD19 (1:80, BD Bioscience, 751213), PacBlue Ly6G (1:250, BD Bioscience, 127611) and PE-Dazzle 594 CD11c (1:500, Biolegend, 117347). Cells and antibody mixture were incubated for 30 minutes. Cells were washed once prior to fixation and permeabilization (BD Cytofix/Cytoperm kit, BDB554714) as per manufacturer’s instructions. Cells were then resuspended in BD Perm/wash buffer with the following antibodies: BV650 IL4 (1:50, BD Bioscience, cat#564004), APC-Fire750 IFNγ (1:80, Biolegend, 505859), AF647 IL-17a (1:200, Biolegend, 506911) and PE-Cy7 Arg1 (1:100, ThermoFisher, 25-3697-80). Cells were incubated with antibody mixture for 30 minutes. Cells were washed twice with BD Perm/wash buffer followed by resuspension in a final volume of 100 μL for flow cytometry analysis.

All samples were analyzed using the Cytek Aurora spectral flow cytometer (Cytek Biosciences, CA, USA). Fluorescence minus one (FMO) samples were used to guide gating strategies shown in Supplementary Fig. 1b. Flow cytometry data was analyzed with the software FCSExpress (DeNovo Software, CA, USA).

### Statistics and reproducibility

Statistical software (GraphPad Prism Version 9.5.1 (528)) was used to analyse data presented as mean with standard deviation (SD). Exact statistical test, p-values and sample sizes are provided in figure legends.

## Data availability

The data supporting the findings of this study are available within the paper and its Supplementary Information.

## Acknowledgements

Funding for this work was provided in part by the James and Kathleen Cornelius Endowment at MSU. J.M. Hix assisted with euthanasia and harvesting tissues. J. Ernst and P. Kubes kindly read and provided feedback on this study.

## Author contributions

Conceptualization, C.V.M. and C.H.C.; Methodology, C.V.M., A.S., A.V.M., E.U., K.B.S., H.P., M.M.K, O.M.B., A.T., M.A., A.S., S.C., A.J.O., K.D.H., R.N., S.P., J.H.E., and C.H.C.; Investigation, C.V.M., A.S., A.V.M., E.U., H.P., M.M.K, O.M.B., A.T., M.A., S.C.; Writing – Original Draft, C.V.M.; Writing – Review & Editing, C.V.M., A.S., A.V.M., E.U., K.B.S., H.P., M.M.K., O.M.H., A.T., M.A., A.K., S.C., A.J.O., K.D.H., R.N., S.P., J.H.E., and C.H.C.; Funding Acquisition, C.H.C.; Resources, R.N. and C.H.C.; Supervision, A.J.O., K.D.H., R.N., S.P., J.H.E., and C.H.C.

---

**Supplementary Figure 1.**
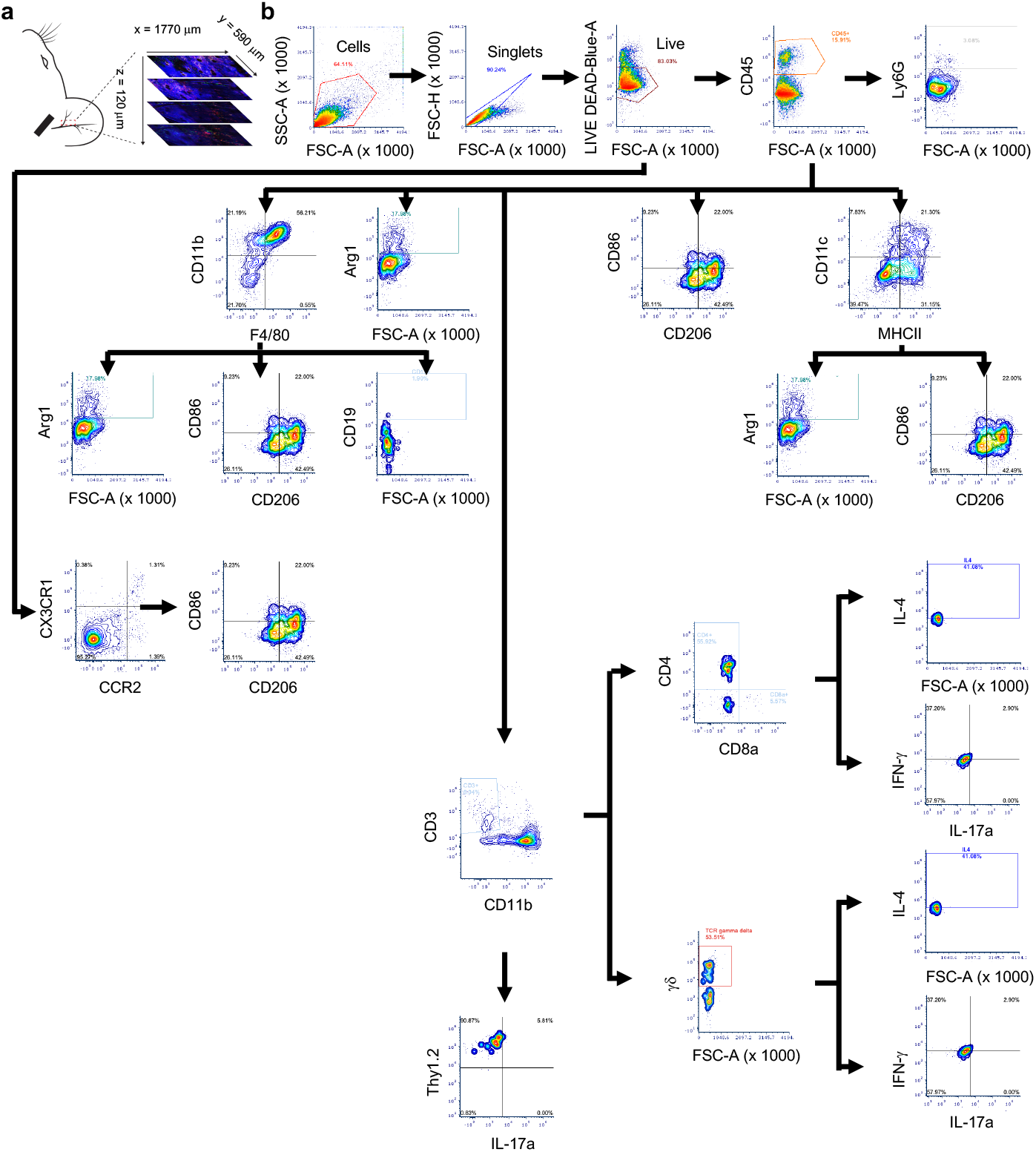
Schematic diagrams showing area imaged by intravital imaging and gating strategy used in flow cytometric analyses. **a**, The base of the mouse ear was surgically incised (sham) and implanted with biomaterials, whereas the adjacent rectangular biomaterial microenvironment was imaged. **b**, Flow chart showing gating strategy, with some data repeated for clarity.

**Supplementary Figure 2.**
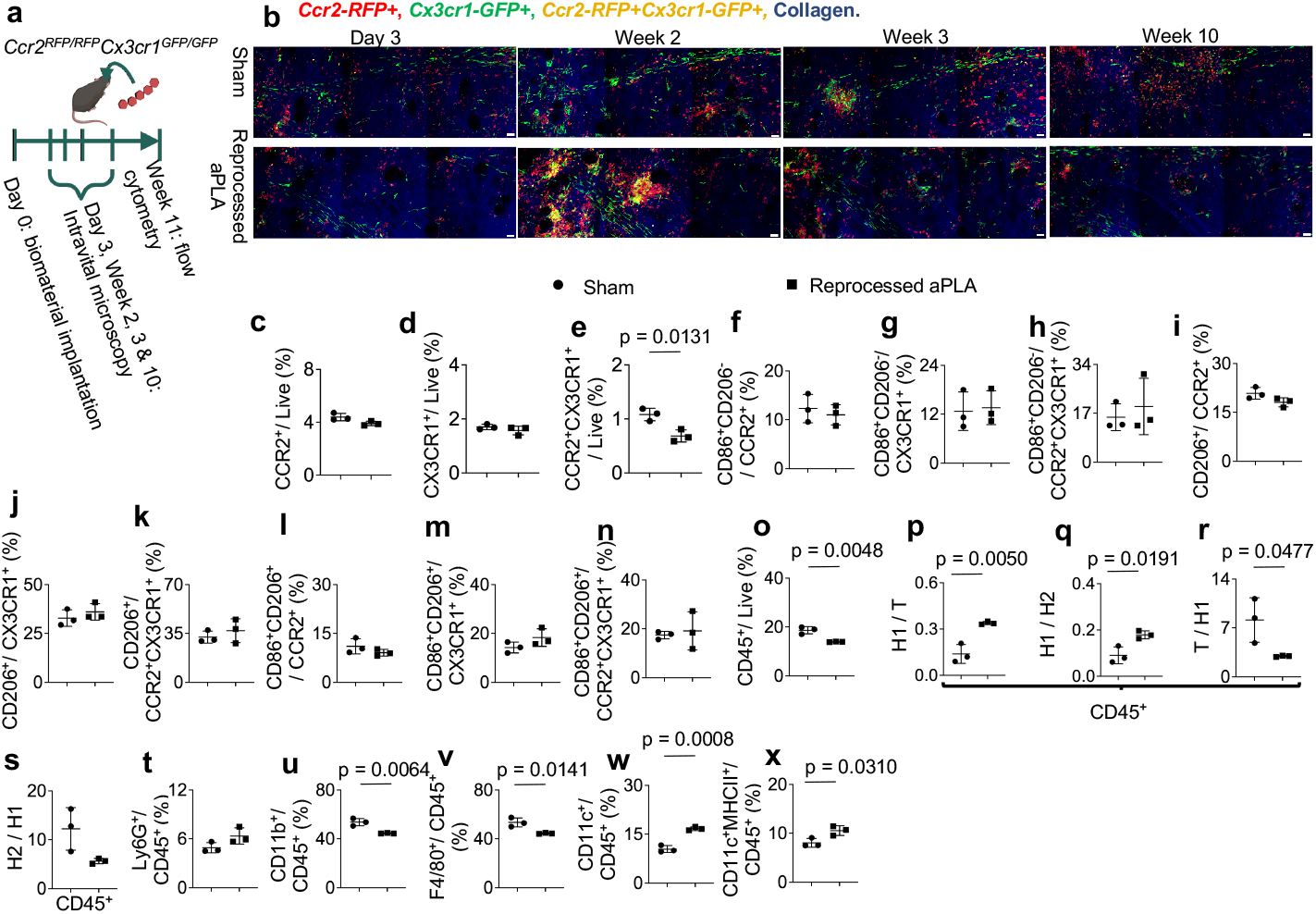
Deficiency of CCR2 and CX3CR1 differentially affects the proportion and activation states of cells in the amorphous polylactide biomaterial microenvironment. **a**, *Ccr2*^*RFP/RFP*^*Cx3cr1*^*GFP/GFP*^ (CCR2- and CX3CR1-deficient) mice were surgically incised (sham group) or implanted with reprocessed amorphous polylactide (aPLA). Afterwards, intravital microscopy and flow cytometric analysis of tissues around incision sites (sham group) or implants were undertaken. **b**, Representative intravital microscopy images at different time points around incision (sham group) or implants (scale bars, 50 μm). **c-e**, Flow cytometry quantification of CCR2^+^(c), CX3CR1^+^(d) and CCR2^+^CX3CR1^+^(e) cells. **f-h**, Quantification of proinflammatory (CD86^+^CD206^-^) cells among CCR2^+^(f), CX3CR1^+^(g) and CCR2^+^CX3CR1^+^(h) populations. **i-k**, Quantification of anti-inflammatory (CD206^+^) cells among CCR2^+^(i), CX3CR1^+^(j) and CCR2^+^CX3CR1^+^(k) populations. **l-n**, Quantification of transition (CD86^+^CD206^+^) cells among CCR2^+^(l), CX3CR1^+^(m) and CCR2^+^CX3CR1^+^(n) populations. **o**, Nucleated hematopoietic (CD45^+^) cells. **p-q**, Fold change of proinflammatory (H1; CD86^+^CD206^-^) cells with respect to transition (T; CD86^+^CD206^+^) cells (p) or anti-inflammatory (H2; CD206^+^) cells (q). **r-s**, Fold change of T (r) or H2 (s) cells with respect to H1 cells. **t**, Neutrophils (CD45^+^Ly6G^+^ cells). **u**, Monocytes (CD45^+^CD11b^+^ cells). **v**, Macrophages (CD45^+^F4/80^+^ cells). **w**, Dendritic cells (CD45^+^CD11c^+^ cells). **x**, Dendritic cells expressing MHCII (CD45^+^CD11c^+^MHCII^+^ cells). Unpaired t-test (two-tailed), n = 3 mice per group.

**Supplementary Figure 3.**
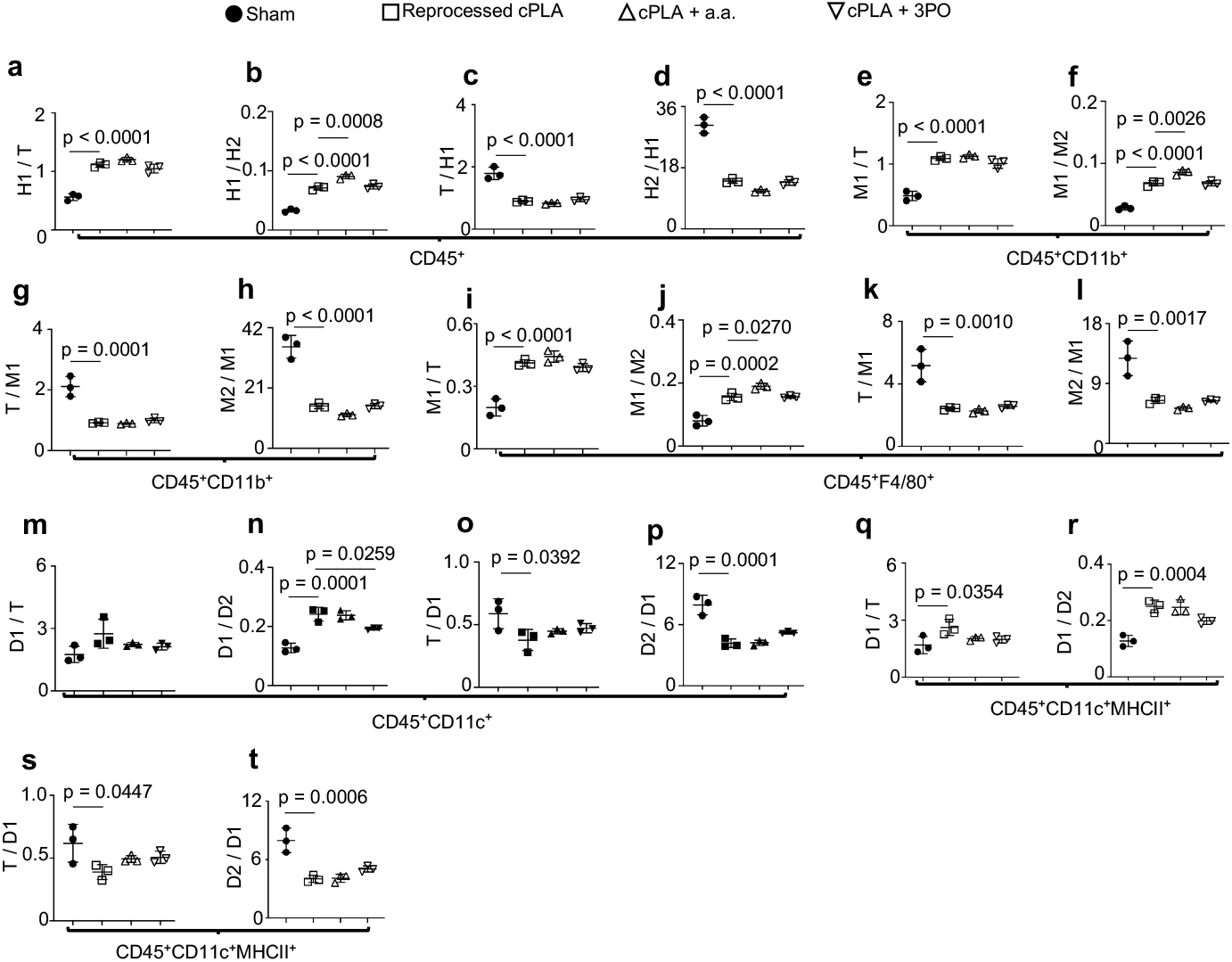
CD86 and CD206 proportions in myeloid populations are not modulated by locally targeting immunometabolism at 11 weeks post-implantation of crystalline polylactide biomaterials. **a-b**, Fold change of proinflammatory (H1; CD86^+^CD206^-^) cells with respect to transition (T; CD86^+^CD206^+^) cells (a) or anti-inflammatory (H2; CD206^+^) cells (b), gated for nucleated hematopoietic (CD45^+^) populations. **c-d**, Fold change of T (c) or H2 (d) cells with respect to H1 cells. **e-f**, Fold change of proinflammatory (M1; CD86^+^CD206^-^) monocytes (CD45^+^CD11b^+^) with respect to transition (T; CD86^+^CD206^+^) monocytes (e) or anti-inflammatory (M2; CD206^+^) monocytes (f). **g-h**, Fold change of T (g) or M2 (h) monocytes with respect to M1 monocytes. **i-j**, Fold change of proinflammatory (M1; CD86^+^CD206^-^) macrophages (CD45^+^F4/80^+^) with respect to transition (T; CD86^+^CD206^+^) macrophages (i) or anti-inflammatory (M2; CD206^+^) macrophages (j). **k-l**, Fold change of T (k) or M2 (l) macrophages with respect to M1 macrophages. **m-n**, Fold change of proinflammatory (D1; CD86^+^CD206^-^) dendritic (CD45^+^CD11c^+^) cells with respect to transition (T; CD86^+^CD206^+^) dendritic cells (m) or anti-inflammatory (D2; CD206^+^) dendritic cells (n). **o-p**, Fold change of T (o) or D2 (p) dendritic cells with respect to D1 dendritic cells. **q-r**, Fold change of proinflammatory (D1; CD86^+^CD206^-^) MHCII^+^ dendritic cells with respect to transition (T; CD86^+^CD206^+^) MHCII^+^ dendritic cells (q) or anti-inflammatory (D2; CD206^+^) MHCII^+^ dendritic cells (r). **s-t**, Fold change of T (s) or D2 (t) MHCII^+^ dendritic cells with respect to D1 MHCII^+^ dendritic cells. One-way ANOVA followed by Tukey’s multiple comparison test, n = 3 mice per group; crystalline polylactide, cPLA; aminooxyacetic acid, a.a.; 3-(3-pyridinyl)-1-(4-pyridinyl)-2-propen-1-one, 3PO.

